# Inhibition of the oxytocin system prevents partner preference formation in females of the monogamous goby *Trimma marinae*: a comparison with its polygamous relative, *Trimma caudomaculatum*

**DOI:** 10.64898/2026.11.26.769516

**Authors:** Kazuya Fukuda, Natsumi Tsujita, Hisato Kuniyoshi, Takao Mukuda, Masayuki Yoshida, Tomoki Sunobe

**Author notes:** Correspondence: Kazuya Fukuda. 1-15-1 Kitazato, Minami-ku, Sagamihara, Kanagawa 252-0373, Japan.

## Abstract

Monogamous mating systems are widespread across vertebrates. Although ecological theories have addressed their ultimate drivers, the extent to which their proximate mechanisms are shared across taxa remains unclear. We investigated the involvement of oxytocin (OT) signaling in partner preference formation in two closely related gobiid fishes with contrasting mating systems: monogamous *Trimma marinae* and polygamous *Trimma caudomaculatum*. Females received pharmacological treatments before cohabitation with a male, and partner preference was assessed the following day. In *T. marinae*, vehicle- and OT-treated females spent significantly more time near the cohabitation male than near a novel male, whereas no significant preference was detected after treatment with atosiban, an OT receptor antagonist. This finding suggests that OT may play an important role in partner preference formation in fishes, as in mammals. On the other hand, administering OT to *T. caudomaculatum* did not induce partner preference, raising the possibility that interspecific differences in neural mechanisms downstream of OT signaling may contribute to the contrasting mating systems of these species. This study adds a finding from fishes to research on the repeated evolution of monogamous mating systems within the framework of partially shared mechanisms.

**Highlights:**

- Oxytocin may contribute to partner preference formation in monogamous fish.
- Oxytocin did not induce partner preference in a polygamous relatives.
- Differences downstream of oxytocin may contribute to contrasting mating systems.
- Findings extend research on partially shared mechanisms in the evolution of monogamy.

## Introduction

Monogamy is a mating system in which males and females typically mate with only one partner during a specified period. It occurs across diverse vertebrate taxa, often accompanied by similar behavioral characteristics (Klug, 2018; Kvarnemo, 2018).

Several ecological hypotheses have been proposed within general ecological frameworks to explain the evolution and adaptive significance of monogamy across taxa (Emlen and Oring, 1977; Klug, 2018; Kvarnemo, 2018). However, research on the proximate mechanisms underlying monogamous behavior and pair bonds, selective and enduring attachments between partners (Bales et al., 2021), has focused primarily on particular taxonomic groups, such as mammals (López-Gutiérrez et al., 2022; Blumenthal and Young, 2023).

Oxytocin (OT), here used as a common name for its vertebrate orthologs including isotocin in teleosts following Theofanopoulou et al. (2021), is a neuropeptide widely conserved across vertebrates and implicated in pair bonding across diverse taxa (Johnson and Young, 2015; Bales et al., 2021; Rappeneau and Castillo Díaz, 2024). Studies in mammals have demonstrated a role for OT in pair bond formation and have revealed its effects on social cue processing and reward-related plasticity (Froemke and Young, 2021; Blumenthal and Young, 2023). Although recent findings in fishes suggest a role for the OT system in mate choice and pair bonding (Yokoi et al., 2020; Nowicki et al., 2020), functional studies in species forming enduring pair bonds remain limited. Extending this research to phylogenetically diverse taxa, including fishes, which are evolutionarily distant from mammals, may identify shared and divergent mechanisms underlying monogamy and provide a more comprehensive understanding of its evolution.

In the present study, we investigated the role of OT in mating systems by comparing two gobiid fishes, *Trimma marinae* and *Trimma caudomaculatum*, which are closely related (Sunobe et al., 2017) and share similar ecologies but differ in mating system.

Although both species inhabit rocky and coral reefs and school near sheltering substrates (Saeki et al., 2005; Fukuda et al., 2017), *T. marinae* forms enduring monogamous pairs (Fukuda et al., 2017), whereas *T. caudomaculatum* is polygamous (Tomatsu et al., 2018). In both species, males defend their nests and generally court approaching females. However, female behavior differs. After engaging in pairing behavior with a male, a female *T. marinae* begins visiting his nest and exhibiting mate guarding behavior that likely contributes to the formation and maintenance of monogamy (Fukuda et al., 2017; Kambe et al., 2026). By contrast, such exclusive pairing and mate guarding have not been reported in females of *T. caudomaculatum* (Tomatsu et al., 2018). Thus, these species provide a useful comparative system for investigating the mechanisms underlying pair bonding.

We therefore administered atosiban, an OT receptor antagonist, to examine whether the OT system is involved in partner preference formation in female *T. marinae*. For comparison, we administered OT to female *T. caudomaculatum* at the start of cohabitation with a male to test whether this treatment induced partner preference. As a preliminary step toward assessing whether our findings could be interpreted in terms of central actions of OT, we also examined the distribution of Evans blue in brain tissue following intraperitoneal administration (Supplementary Material 1).

## Materials and methods

### Animals and housing

Partner preference experiments were conducted from June to September 2016 using 40 males and 20 females of *Trimma marinae* and 22 males and 11 females of *T. caudomaculatum.* Fish were collected in April 2016 using SCUBA and hand nets at 6–24 m depths offshore of Amami Island, Japan, and transported to the laboratory at Tateyama Station, Tokyo University of Marine Science and Technology. Male and female *T. marinae* were housed separately in 182 L tanks for at least one week to prevent pair formation before testing. Because *T. caudomaculatum* undergoes socially regulated bidirectional sex change depending on body size (Tomatsu et al., 2018), both sexes were maintained together in a 243 L tank. Females were transferred in groups of six to 65 L tanks one week before testing. Additional large females beyond the 11 focal females were maintained so that the largest female in each tank could be excluded from the experiment without altering the social holding conditions. Tanks were maintained at 25–26 °C under the natural photoperiod, and fish were fed twice daily.

### Drug administration

Oxytocin (OT; isotocin, [Ser^4^,Ile^8^]-oxytocin; Bachem, California, USA) and atosiban (a peptide OT/vasopressin receptor antagonist; Sigma-Aldrich, St. Louis, MO, USA) were dissolved in 0.9% NaCl. To minimize handling stress before testing, injection concentrations were calculated using the mean body masses of a reference sample of females collected at the same time and maintained under the same conditions (*T. marinae*: mean ± SD = 0.18 ± 0.02 g, n = 30; *T. caudomaculatum*: 0.52 ± 0.02 g, n = 20). Females received a dose of approximately 10 μg g ¹ body weight in 0.5 μl (*T. marinae*) or 1 μl (*T. caudomaculatum*). For anesthesia, 2-methylquinoline was mixed with 99% ethanol at a 1:1 ratio (v/v), and the resulting mixture was diluted in seawater to a final concentration of 18 ppm. Females were anesthetized in this solution and injected intraperitoneally using a 5 μl Hamilton syringe with a 33-gauge needle.

### Partner preference test

The experimental sequence is shown schematically in Fig. 1. On day 1, a focal female was injected with OT, atosiban, or vehicle and immediately placed with a randomly selected male (hereafter, the cohabitation male) in a 60 × 30 × 30 cm tank containing coral sand and a half-pipe spawning shelter. Fish cohabited for 6 h during the species-specific reproductive period (05:00–11:00 for *T. marinae* and 11:00–17:00 for *T. caudomaculatum*). For both species, only pairs in which the male actively courted the female and the female remained near the male, but spawning did not occur, were retained for the subsequent partner preference test. After cohabitation, the female and cohabitation male were housed in separate, isolated tanks until the following day. On day 2, the female was tested in a 60 × 30 × 30 cm tank divided into one female compartment and two adjacent male compartments. The cohabitation male and a novel male matched to within 0.5 mm in total length were simultaneously visible to the focal female through transparent partitions but not to each other. To prevent potential side bias from affecting the results, the left–right positions of the cohabitation and novel males were randomly assigned for each trial. Since the partitions were not sealed to the tank, there was a slight water exchange between the compartments. After 30 min of acclimation in a centrally positioned opaque 2 L chamber in the female compartment, the female was released and recorded from above the test tank for 30 min. During the test, we confirmed whether both males actively displayed courtship behavior. A researcher not involved in behavioral scoring randomized the order of the videos, and the observer was blind to treatment. The observer measured the association time (s) during which the female’s snout was within 5 cm of each male’s compartment. After excluding trials in which one or both stimulus males failed to display courtship during the preference test, data from 18 *T. marinae* females (OT, n = 6; atosiban, n = 5; vehicle, n = 7) and 11 *T. caudomaculatum* females (OT, n = 6; vehicle, n = 5) were included in the analysis.

**Fig. 1.**
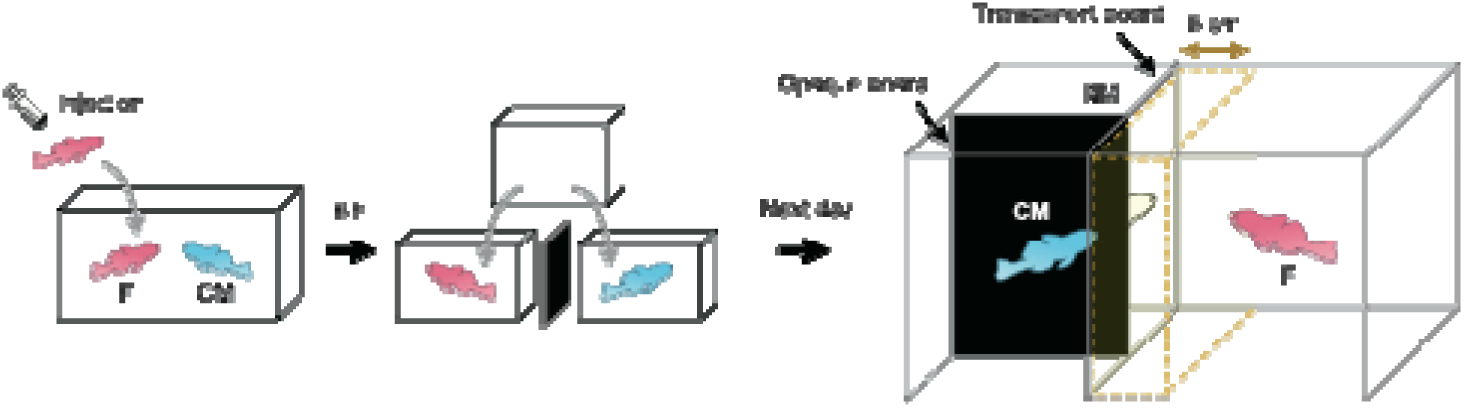
Schematic overview of the partner preference test. On day 1, a focal female was injected with OT, atosiban, or vehicle and allowed to cohabit with a male (cohabitation male) for 6 h. The female and cohabitation male were then housed separately in isolated tanks until the following day. On day 2, the female was simultaneously presented with the cohabitation male and a novel male in adjacent compartments. Association time was defined as the time during which the female’s snout was within 5 cm of each male’s compartment during the 30 min test. *Pink*, focal female (*F*); *light blue*, cohabitation male (*CM*); *white*, novel male (*NM*)

### Statistical analyses

Behavioral data are presented as means ± standard errors of the mean. Reference body masses are presented as means ± SD. Analyses were performed in R version 4.4.1 (R Core Team, 2024). The significance level was set at α = 0.05. Within each species–treatment group, association times with the cohabitation male (*C*) and novel male (*N*) were compared using a paired *t*-test, with Hedges’ *g_z_* as the effect size. Partner preference scores were calculated as (100*C*/(*C* + *N*)). Scores were compared among the vehicle, OT, and atosiban groups in *T. marinae* using Welch’s one-way ANOVA with approximate partial η *²* as the effect size, and between the OT and vehicle groups in *T. caudomaculatum* using Welch’s t-test with Hedges’ *g*. Total association time was calculated as (*C*+*N*) and analyzed using the same between-treatment procedures.

### Ethical statement

At Tokyo University of Marine Science and Technology, there were no institutional ethical regulations specifically for research on fish at the time this study was conducted. Therefore, all procedures followed the Guidelines for the treatment of animals in behavioural research and teaching (ASAB/ABS, 2012).

## Results

### Effect of pharmacological treatment on partner preference in T. marinae

Females treated with vehicle, OT, or atosiban spent 1388.4 ± 94.7 s, 1241.7 ± 176.7 s, and 885.8 ± 321.8 s, respectively, near the cohabitation male, and 176.3 ± 78.2 s, 185.0 ± 122.3 s, and 322.6 ± 167.5 s, respectively, near the novel male. Association times differed significantly between the cohabitation and novel males in the vehicle group (*t* = 8.02, *p* < 0.001, *g_z_* = 2.63) and OT group (*t* = 3.80, *p* = 0.013, *g_z_* = 1.30), with females in both groups spending more time near the cohabitation male. No significant difference was detected in the atosiban group (*t* = 1.32, *p* = 0.258, *g_z_* = 0.47; Fig. 2A). Mean partner preference scores were 89.2 ± 4.7%, 86.1 ± 8.8%, and 73.5 ± 14.4% in the vehicle, OT, and atosiban groups, respectively. There was no significant difference among treatments (*F*, = 0.506, *p* = 0.622, η*_p_*² = 0.118; Fig. 2C). Total association times were 1564.7 ± 85.7 s, 1426.7 ± 122.4 s, and 1208.4 ± 283.6 s in the vehicle, OT, and atosiban groups, respectively, and did not differ significantly among treatments (*F*,. = 0.915, *p* = 0.439, η*_p_*² = 0.189; Fig. 2D).

**Fig. 2.**
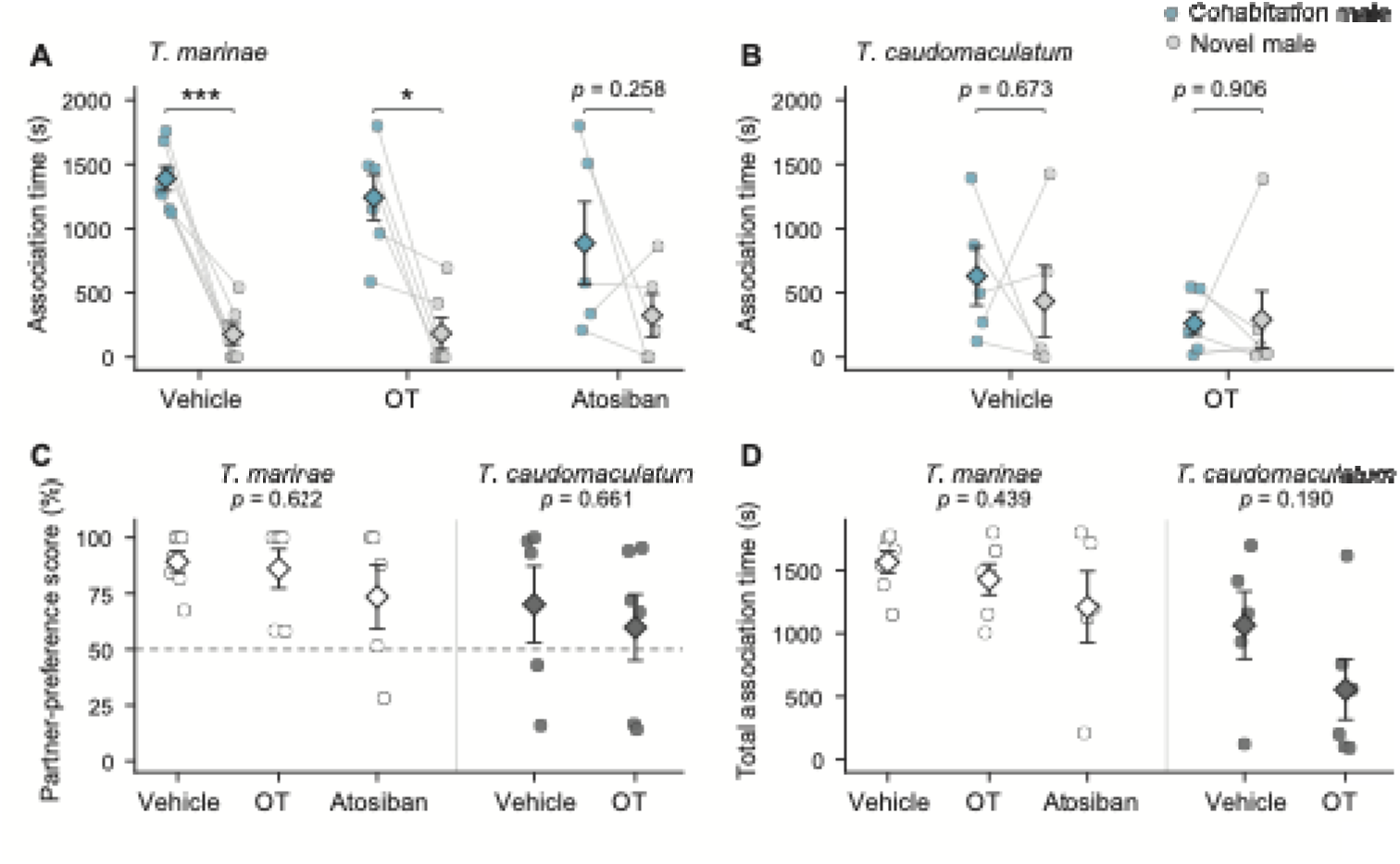
Effects of pharmacological treatment on partner preference in female *Trimma marinae* and *T. caudomaculatum*. Association times with the cohabitation male (*blue*) and novel male (*light-grey*) are shown for *T. marinae* (**A**) and *T. caudomaculatum* (**B**). *Circles* represent each female with values from the same female connected by grey lines. *Diamonds* and error bars indicate means ± SEM. Partner preference scores (**C**). The *dashed horizontal line* indicates the score expected in the absence of preference (50%). Total association time (**D**). In C and D, *white symbols* represent *T. marinae* and *dark-grey symbols* represent *T. caudomaculatum*. *T. marinae*, vehicle, n = 7; OT, n = 6; atosiban, n = 5; *T. caudomaculatum,* vehicle, n = 5; OT, n = 6. Within-group comparisons in A and B used paired *t*-tests. Between-treatment comparisons in C and D used one-way ANOVA for *T. marinae* and Welch’s *t*-tests for *T. caudomaculatum*. *p < 0.05, ***p < 0.001

### Effect of OT on partner preference in female T. caudomaculatum

Females treated with vehicle or OT spent 630.2 ± 228.3 s and 262.3 ± 92.9 s, respectively, near the cohabitation male, and 434.6 ± 276.5 s and 292.2 ± 220.8 s, respectively, near the novel male. Association times with the cohabitation and novel males did not differ significantly in either the vehicle group (*t* = 0.45, *p* = 0.673, *g_z_* = 0.16) or the OT group (*t* = −0.12, *p* = 0.906, *g_z_* = −0.04; Fig. 2B). Mean partner preference scores were 70.1 ± 17.2% in the vehicle group and 59.8 ± 14.8% in the OT group, with no significant difference between treatments (*t*. = −0.45, *p* = 0.661, *g* = −0.25; Fig. 2C). Total association times were 1064.8 ± 267.6 s and 554.5 ± 238.5 s in the vehicle and OT groups, respectively, and also did not differ significantly between treatments (*t*. = −1.42, *p* = 0.190, *g* = −0.79; Fig. 2D).

## Discussion

Our findings suggest a role for OT signaling in partner preference formation in female *T. marinae*. OT may play an important role in the formation of enduring pair bonds in this species, as in mammals, although a contribution of vasotocin receptor inhibition cannot be excluded because atosiban also antagonizes vasopressin receptors. The findings also raise the possibility that the results could be interpreted in terms of central OT actions (Supplementary Material 1). However, partner preference scores did not differ significantly among treatments. Further comparative studies should determine how much OT contributes to partner preference formation in fishes and whether its contribution varies across species and social contexts.

OT administration did not induce partner preference in *T. caudomaculatum,* raising the possibility that interspecific differences in mechanisms downstream of OT (e.g., receptor expression patterns and the processing mechanisms linking ligand reception to monogamous behavior) contribute to the establishment of contrasting mating systems in the two species. This hypothesis is consistent with evidence linking receptor expression to partner preference in prairie voles (Blumenthal and Young, 2023). These parallels suggest that partially shared neuroendocrine mechanisms may contribute to the repeated evolution of monogamy across vertebrates (Fischer et al., 2019). Comparing how shared systems generate partner preference through neural circuits across species may advance our understanding of the evolution and diversification of monogamy.

## CRediT authorship contribution statement

**Kazuya Fukuda**: Conceptualization, Methodology, Investigation, Formal analysis, Visualization, Resources, Funding acquisition, Project administration, Writing – original draft, Writing – review & editing. **Natsumi Tsujita**: Conceptualization, Writing – review & editing. **Hisato Kuniyoshi**: Conceptualization, Methodology, Writing – review & editing, Supervision. **Takao Mukuda**: Methodology, Writing – review & editing, Supervision. **Masayuki Yoshida**: Methodology, Writing – review & editing. **Tomoki Sunobe**: Conceptualization, Resources, Funding acquisition, Writing – review & editing, Supervision.

## Declaration of generative AI and AI-assisted technologies in the manuscript preparation process

During manuscript preparation, the authors received partial assistance from ChatGPT and Codex (OpenAI) with English-language drafting and revision, and adjustments to the layout and formatting of Fig. 2 and Supplementary Material 2. All statistical analyses were performed by the authors. The authors checked and revised the resulting material and take full responsibility for the final manuscript.

## Data availability

The behavioral data supporting the findings of this study are provided in Supplementary Material 2.

## Acknowledgments

We are grateful to S. Yokoyama for assistance with fish collection. We also thank Dr. N. Yamamoto and Dr. H. Abe for their valuable advice and support.

---

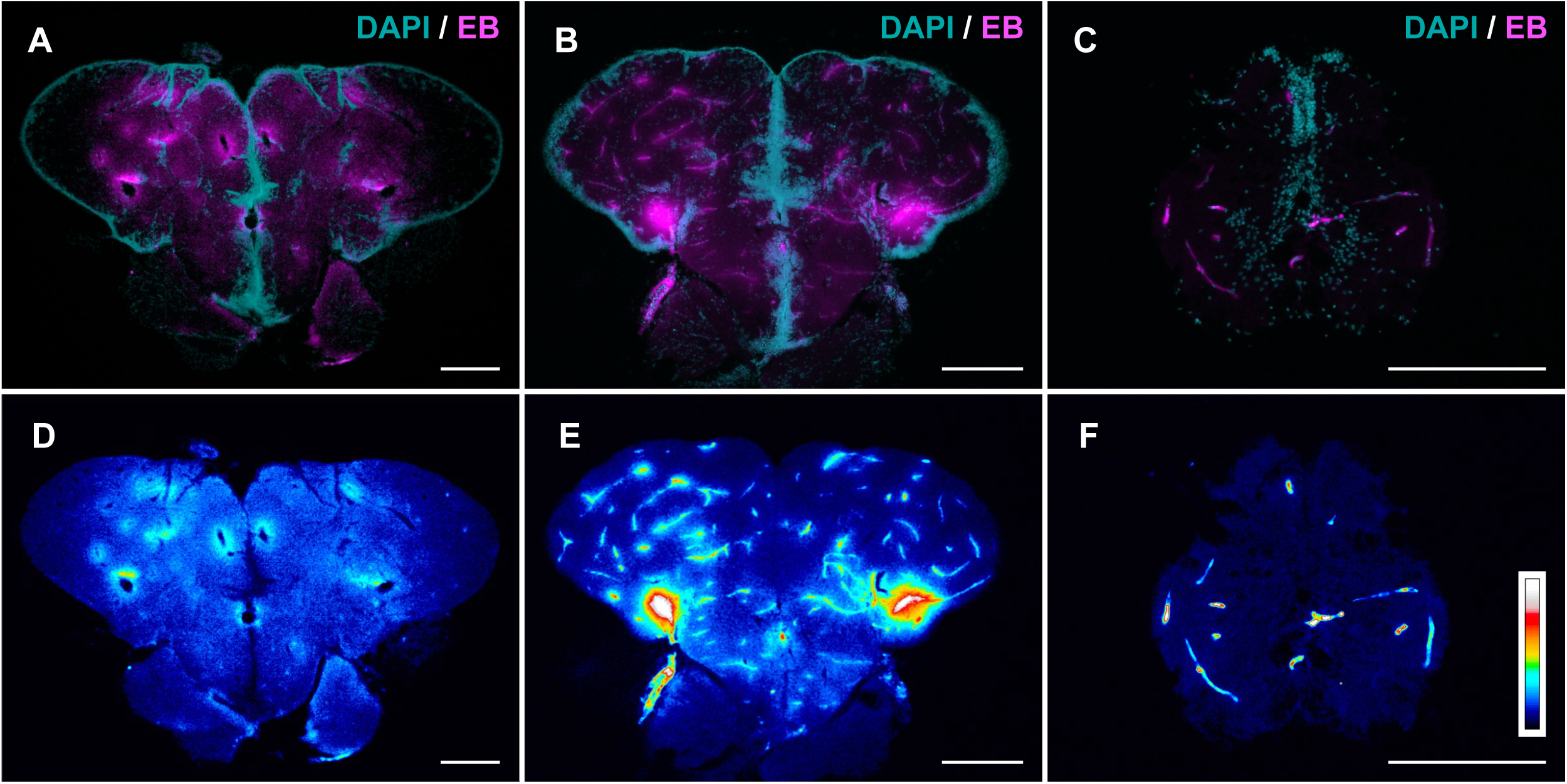

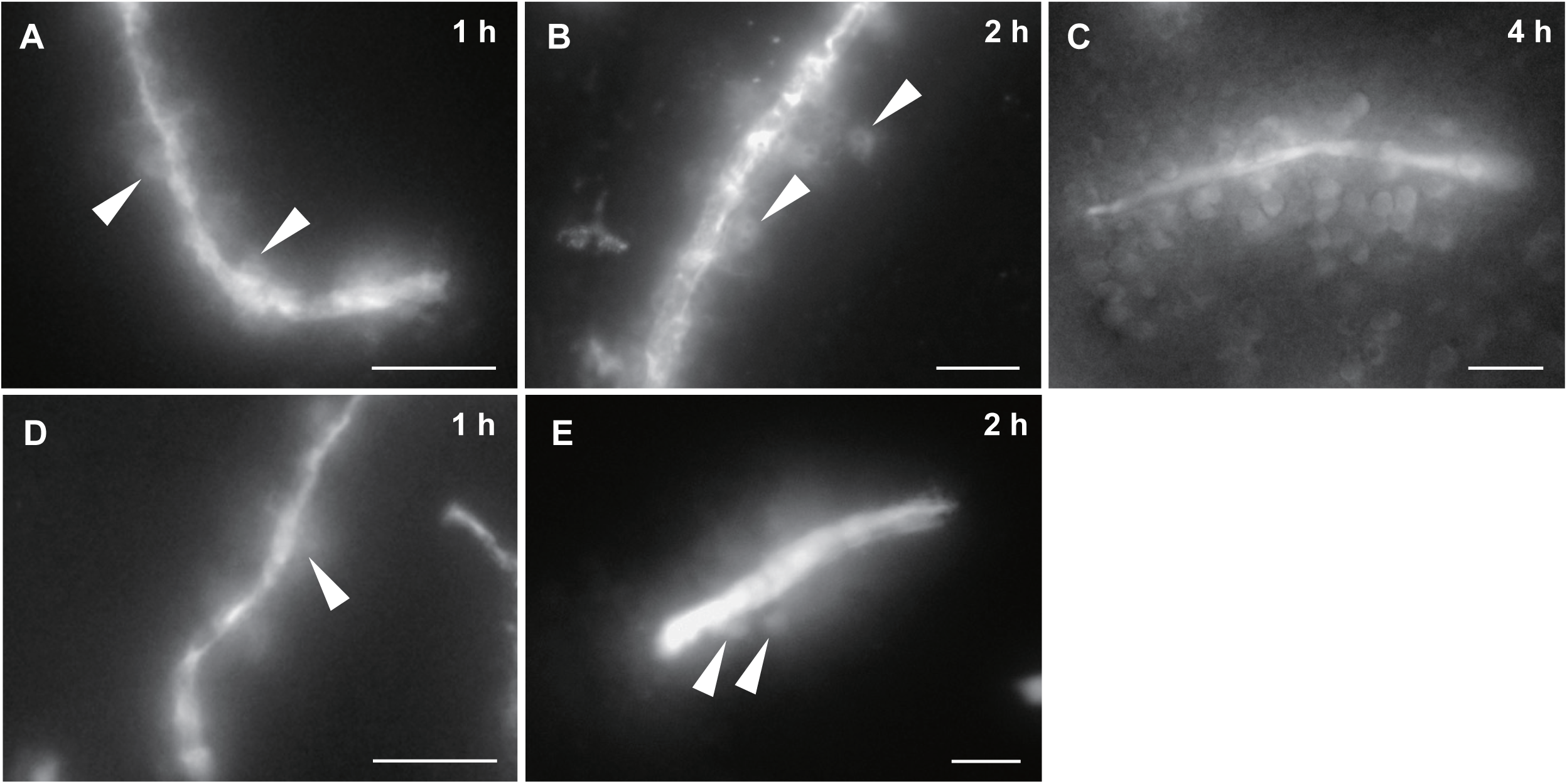

