## Supplementary Material 1 for "Inhibition of the oxytocin system prevents partner preference formation in females of the monogamous goby *Trimma marinae*: a comparison with its polygamous relative, *Trimma caudomaculatum*"

^3^Current address: School of Marine Biosciences, Kitasato University, Sagamihara, Kanagawa 252-0373, Japan

^6^Current address: Department of Zoology, National Museum of Nature and Science, Amakubo, Tsukuba, Ibaraki 305-0005, Japan

**Supplementary methods**

*Animals*

Evans blue (EB) administration was conducted from 2018 to 2020 using eight females each from *T. marinae* and *T. caudomaculatum*. These individuals were collected using the same procedure as described in the main text, and were housed in a 375 L tank at Nagoya University under a 12-h light and 12-h dark photoperiod. Other husbandry conditions were as described in the main text, except that no coral sand was used.

*Evans blue administration and tissue collection*

Fish were anesthetized with ethyl 3-aminobenzoate methanesulfonate (MS-222, Sigma, St. Louis, USA) and injected intraperitoneally with 0.2 µl and 0.5 µl of EB solution, dissolved in phosphate-buffered saline (50 mg/ml EB in PBS [pH 7.4]), into *T. marinae* and *T. caudomaculatum*, respectively. After intervals of 1 or 2 h (n = 3 per species at each interval), the fish were deeply anesthetized with MS-222 and immediately decapitated. When EB is present at a high concentration within the blood vessels, the vessels emit strong fluorescence, which may create the misleading appearance that fluorescence extends into the surrounding brain parenchyma because of the spread of light from the blood vessels. To address this potential artifact, in the case of *T. caudomaculatum* (n = 2), which is relatively large and more accessible to manipulate surgically, we injected EB using the same method as above, allowed a 4 h interval, then performed transcardial perfusion fixation under deep anesthesia to flush out the blood from the vessels. In contrast, for *T. marinae* (n = 2), which are smaller and challenging to manipulate surgically, we decapitated the fish 4 h after EB injection and immersed the brain in a fixative solution (4% paraformaldehyde in 0.1 M PB [pH 7.4]). For comparison, we also observed spinal cord sections from the same specimens.

*Fixation and section preparation*

For fish that were not perfused, the brains were surgically removed from the skulls in fixative (4% paraformaldehyde in 0.1 M PB [pH 7.4]) pre-chilled on ice. The brains were post-fixed in the same solution at 4 °C for 2 h in the dark. From this point onward, all procedures were carried out under low-light conditions. The fixed brains were cryoprotected by immersion in 20% sucrose in 0.1 M PB at 4 °C until the brains sank in the solution, approximately 4 h. They were embedded in 5% agarose type IX-A (Sigma, St. Louis, USA) in 20% sucrose in 0.1 M PB and frozen in n-hexane at approximately -60 °C. Transverse sections were cut at a thickness of 16 µm on a cryostat and mounted on gelatin-coated glass slides. Spinal cord samples were processed into sections and mounted on slides following the same procedure.

*Observation of unwashed sections*

Since EB is a water-soluble dye, washing the sections with PBS can weaken the fluorescence signal. Therefore, sections from fish sampled 1 or 2 h after EB injection were air-dried in the dark at room temperature for about 10 min and observed without washing or further processing under a fluorescence microscope (BX60, Olympus, Tokyo, Japan). Since EB has been reported to be taken up by neurons through endocytosis and to accumulate in their cell bodies (Hamasaki et al., 2020), we examined fluorescence associated with cell-body-like structures around blood vessels in the telencephalon at different post-injection intervals. In this study, we focused on the telencephalon, which contains regions involved in various social behaviors in teleost fishes (O’Connell and Hofmann, 2011).

*DAPI staining and image processing*

Only sections from the 4 h experiments (n = 2 per species) were processed for DAPI staining after sectioning. The sections were air-dried in the dark at room temperature for approximately 60 min, then washed twice in PBS for 3 min each. The sections were incubated in PBS containing DAPI at 1 µg/ml for 10 min at room temperature, followed by three washes in PBS for 3 min each, and were then mounted using Fluoromount (Diagnostic BioSystems, California, USA). Fluorescence microscopy and a digital camera were used to photograph the sections. In some images, pseudo-coloring was applied using the “royal” lookup table in ImageJ to enhance the visual differentiation of EB fluorescence intensity based on brightness.

*Ethical statement*

All procedures followed the Nagoya University Regulations on Animal Care and Use in Research, and Guidelines for the treatment of animals in behavioural research and teaching (ASAB/ABS, 2012).

**Supplementary results**

In both *T. caudomaculatum* and *T. marinae* examined 4 h after EB injection, EB-associated fluorescence was observed around blood vessels in the telencephalon (Fig. S1A, B, D, E). These perivascular patterns were observed in all individuals examined at this interval. In *T. caudomaculatum*, fluorescence was detected in the brain parenchyma surrounding blood vessels following perfusion (Fig. S1A, D). Similarly, in *T. marinae*, fluorescence extended into the surrounding brain parenchyma (Fig. S1B, E). In spinal cord sections from the same *T. marinae* individuals, fluorescence was mainly associated with blood vessels, with no clearly discernible perivascular spread (Fig. S1C, F).

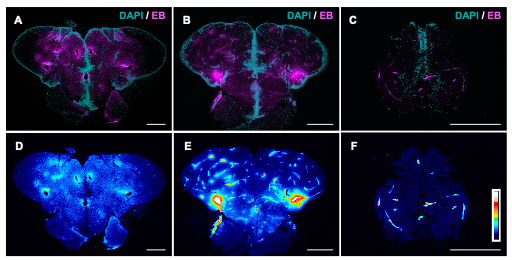

**Figure S1. Distribution of Evans blue-associated fluorescence in the telencephalon and spinal cord.** Representative transverse sections obtained 4 h after intraperitoneal Evans blue (EB) injection. Telencephalon of *T. caudomaculatum* following perfusion (**A**, **D**) and *T. marinae* without perfusion (**B**, **E**). Panels **C** and **F** show the spinal cord of the same *T. marinae* individual shown in **B** and **E**. All sections were imaged after PBS washing, DAPI staining, and mounting. (**A–C**) Merged images showing DAPI in cyan and EB-associated fluorescence in magenta. (**D–F**) The corresponding EB channels displayed using the “royal” lookup table in ImageJ. The color key in **F** indicates the display color sequence from low (black) to high (white) fluorescence intensity. The correspondence between fluorescence intensity and color was not standardized across images; the color key therefore does not provide a common quantitative intensity scale between panels. Scale bars, 200 µm.

Detailed observation of the perivascular brain parenchyma in the telencephalon revealed fluorescence associated with cell-body-like structures in both *T. marinae* and *T. caudomaculatum* (Fig. S2). Although exact quantification was not conducted in this study, these fluorescent structures appeared more numerous at longer intervals after EB injection. In Fig. S2, the 4 h time point is represented only by a non-perfused *T. marinae* specimen (Fig. S2C). EB has been reported to be taken up by living neurons through endocytosis and to accumulate in their cell bodies, making small amounts of dye more readily detectable by fluorescence microscopy (Hamasaki et al., 2020). Thus, the fluorescence associated with cell-body-like structures may reflect cellular uptake of EB following its extravasation into the surrounding brain parenchyma.

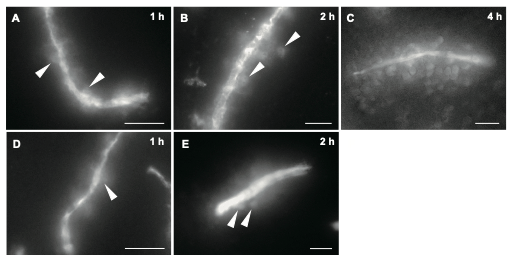

**Figure S2. Evans blue-associated fluorescence around blood vessels in the telencephalon.** Representative images showing EB-associated fluorescence in grayscale. (A–C) *T. marinae* examined 1, 2, and 4 h after EB injection, respectively. (D, E) *T. caudomaculatum* examined 1 and 2 h after EB injection, respectively. All *T. caudomaculatum* specimens examined at 4 h underwent perfusion. Therefore, no images from non-perfused specimens were obtained at this time point. Arrowheads in A, B, D, and E indicate examples of fluorescence associated with cell-body-like structures around blood vessels. All specimens shown were immersion-fixed without perfusion. Although sections in A, B, D, and E were imaged without washing or DAPI staining, the section in C was imaged after PBS washing, DAPI staining, and mounting. Only the EB channel is displayed. Scale bars, 20 µm.
